# Gemcitabine plus nab-paclitaxel preserves skeletal and cardiac mass and function in a murine model of pancreatic cancer cachexia

**DOI:** 10.1101/2023.04.15.536434

**Authors:** Ashok Narasimhan, Daenique H.A Jengelley, Joshua R. Huot, Tara S. Umberger, Emma H. Doud, Amber L Mosley, Meijing Wang, Xiaoling Zhong, Brittany R. Counts, Joseph E. Rupert, Andrew R Young, Andrea Bonetto, Daniel J Horan, Alexander G. Robling, Melissa L. Fishel, Mark R. Kelley, Leonidas G. Koniaris, Teresa A. Zimmers

**Author notes:** Correspondence: Teresa A. Zimmers, PhD, 2730 S. Moody Ave, Portland, OR 97201.

## Abstract

More than 85% of patients with pancreatic ductal adenocarcinoma (PDAC) suffer from cachexia, a debilitating syndrome characterized by the loss of muscle and fat and remains an unmet medical need. While chemotherapy remains an effective treatment option, it can also induce weight and muscle loss in patients with cancer. Gemcitabine combined with nab paclitaxel (GnP) is a first line treatment option for patients with PDAC but GnP’s effect on cachexia has not been comprehensively investigated. We interrogated the effects of GnP in a murine model of pancreatic cancer cachexia. Mice were orthotopically implanted with the cachexia inducing pancreatic cell line (KPC) and were administered GnP or vehicle. The controls underwent sham surgery. We defined GnP effects on cachexia and tumor burden by evaluating muscle and cardiac mass and function, fat mass, bone morphometry, and hematology measurements. We completed RNA sequencing and deep proteome profiling in skeletal and cardiac muscle. KPC+GnP reduced tumor burden over 50% and increased survival compared to KPC. KPC vehicle group had more than 15% muscle mass loss and decreased left ventricular mass, this was not present in KPC+GnP when compared to controls. RNA Seq and deep proteomics analyses suggested that muscle and cardiac dysfunction pathways activated in KPC group were either reversed or decreased in KPC+GnP. In all, our data suggests that GnP protects against muscle and cardiac wasting in an experimental model of PDAC cachexia.

**Graphical Abstract.**
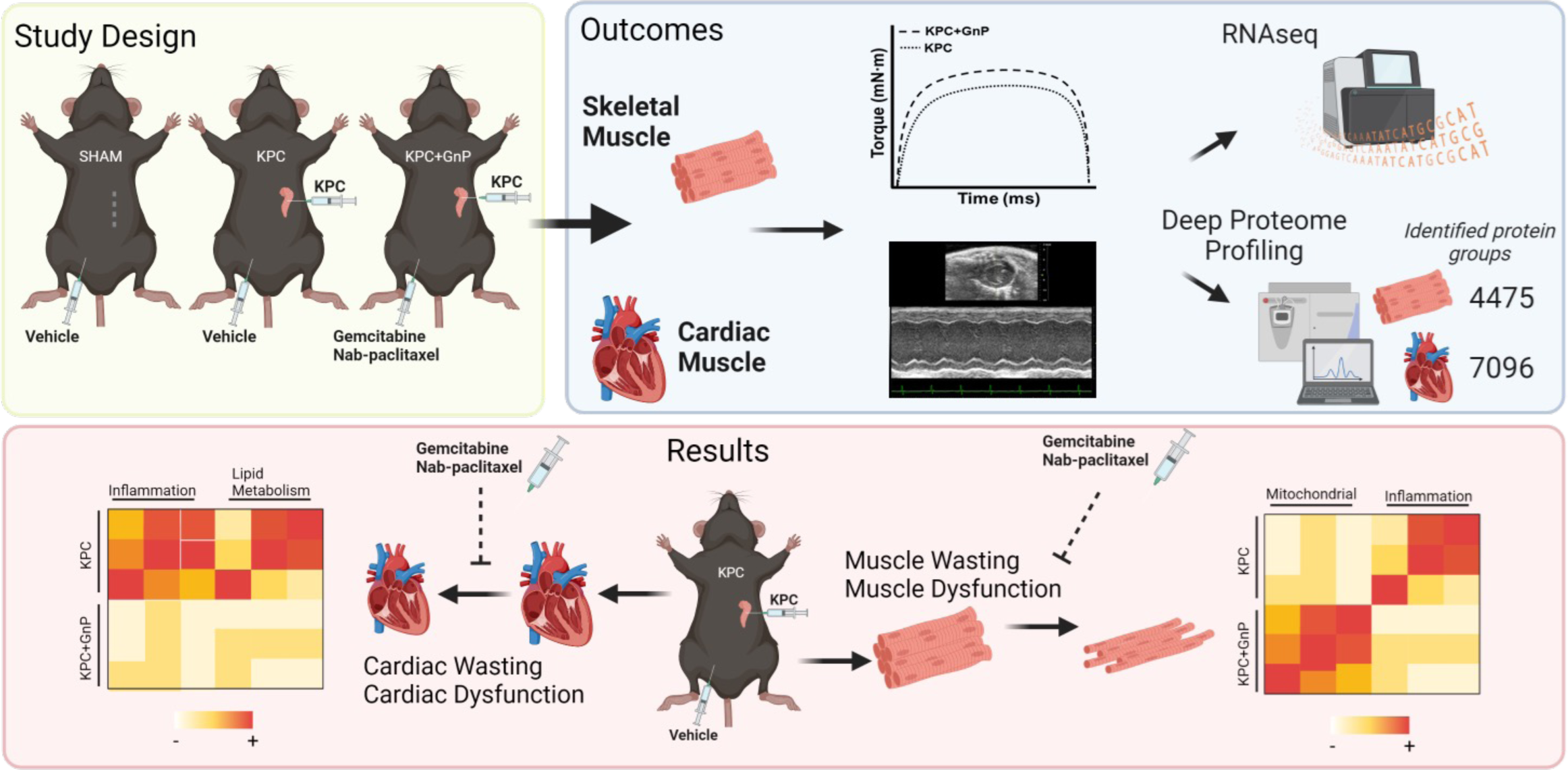

## Introduction

More than 80% of patients with pancreatic ductal adenocarcinoma (PDAC) experience cachexia, a debilitating syndrome characterized by severe muscle and fat wasting^1, 2^. More than 70% of newly diagnosed PDAC cases present with cachexia and it continues to remain as a considerable health burden as cachexia reduces tolerance to therapy^3^ and severely impacts life quality^4^. As there are no approved therapies for cachexia to date, cachexia continues to remain as an unmet medical need requiring more efforts to develop therapeutic interventions.

Patients with cancer are often treated with chemotherapy either in an adjuvant or neo-adjuvant setting. It is well documented that chemotherapy often results in mild to deleterious side effects including nausea, vomiting and anorexia to name a few^5, 6^. Studies have also shown that anticancer agents are beneficial in reducing tumor size. However, several chemotherapeutic agents have a negative impact causing muscle loss and weakness thereby adversely impacting the quality of life^4, 7^. In experimental models, chemotherapy alone (absence of a tumor) induced muscle wasting^8, 9^. In the presence of cancer, cisplatin treatment in mice bearing colon C-26 adenocarcinoma reduced tumor mass but muscle wasting still persisted^4^. Similarly, when C-26 adenocarcinoma mice were treated with Folfiri, the systemic effects continued to persist in the form of muscle wasting and weakness^8^. These studies clearly show that chemotherapy may be detrimental and may aid in cachexia progression. In line with these observations, a comprehensive evaluation on understanding the effect of chemotherapy in PDAC cachexia is currently lacking.

Gemcitabine combined with nab paclitaxel (GnP) is an FDA approved first-line treatment option for patients with locally advanced or metastatic PDAC^10, 11^. Several studies have demonstrated that GnP administration significantly improves the median survival in patients with pancreatic cancer^6, 12^. Prior meta-analysis depicts GnP to be more tolerable in metastatic pancreatic cancer than FOLFIRINOX^13^, yet recent clinical evidence shows FOLFIRIONX increases survival (2 months) and reduce adverse events compared to GnP^14^. Recent work has identified that patients with unresectable pancreatic cancer receiving GnP that exhibited early declines in skeletal muscle had reduced progression free survival^15^. To extend our understanding of GnP’s impact on cachexia, we utilized an experimental model of PDAC cachexia to understand the systemic effects of GnP in skeletal and cardiac muscle, bone morphology and hematology analysis. Using the latest technologies and available interactive software, we are the first to complete deep mass spectrometry proteome profiling in both skeletal and cardiac muscle from cachectic mice. Our results demonstrate that gemcitabine and nab-paclitaxel treatment reduced tumor burden, prevented muscle and cardiac wasting and function, and may potentially be considered as an effective treatment option for pancreatic cancer cachexia.

## Methods

### Cell Culture

KPC 32908 (kindly provided by Dr. David Tuveson) cells were thawed at 37°C in a water bath and grown using DMEM growth medium (10013CV; Corning, NY, USA) supplemented with 10% FBS (SH30071.03, GE Healthcare Life Sciences; Pittsburgh, PA, USA) and 1% pen-strep (15140122, ThermoFisher MA, USA;). In preparation for surgery, the cells were trypsinized when they reached 80% confluency and were counted using a hemocytometer and the required number of cells were suspended in growth media.

### Mice and orthotopic implantation of tumor

12-week-old male C57BL/6J mice were used (000664, Jackson Laboratory; Bar Harbor, Maine, USA). The mice were maintained in 12-hour light and dark cycles and were provided with water *ab-libitum* and allowed to acclimate for a week. The study was done under an approved protocol and the experiments were performed in accordance with the Indiana University School of Medicine Institutional Animal Care and Use Committee.

The surgical procedure for orthotopic implantation of pancreatic cancer cells have been described in detail elsewhere^16^. Briefly, mice were anesthetized using 4% isoflurane and were given a subcutaneous injection of extended-release buprenorphine at 0.5 mg/kg. Once the mice reached the surgical plane, an incision was made in the abdomen retracting the spleen and pancreas. 50,000 KPC 32908 cells prepared in 50uL culture media were injected into the pancreas. After injection, the spleen and pancreas were gently moved inside, and the incision was closed by suture, then the skin was closed using a sterile wound clip. Sham surgery mice underwent the same above procedure without manipulation of the spleen or pancreas.

We repeated the experiment (n=8-10 mice per group) in two different instances. Given no statistical difference in the results, the data was combined from the studies and statistical analysis was conducted on the compiled data.

### Experimental groups and gemcitabine and nab-paclitaxel intervention

The experiment had three groups: sham surgery (Control), KPC group receiving saline as vehicle (KPC) and KPC group receiving gemcitabine and nab-paclitaxel (KPC+GnP). Mice received 120 mg/kg of gemcitabine and 10 mg/kg of nab-paclitaxel on Day 4 and Day 10 after surgery. Gemcitabine and nab-paclitaxel were generously given to us from the Indiana University Health Simon Cancer Center Infusion Center. Injections were administered by the In Vivo Therapeutics Core at IU Simon Comprehensive Cancer Center. Mice included in the analysis were euthanized on day 14. A survival study was completed. Survival study endpoints include less than 5% of total fat mass by EchoMRI, greater than 10% body weight loss or a score of 2 or less on the Hickman body condition score.

### In vivo muscle contractility

To understand the effect of GnP on muscle function, plantarflexion torque assessment was performed (n=10 per group) as previously described (Aurora Scientific Inc., Canada)^17, 18^. Briefly, the left foot was taped and positioned with the foot and tibia positioned at 90 degrees with the knee clamped at the femoral condyles, avoiding fibular nerve compression. The mononuclear electrodes (Natus Neurology, Middleton, WI, USA) were positioned subcutaneously posterior to the knee for tibial nerve stimulation. For determining the maximal stimulus density, peak twitch torque was established. For the assessment of force-frequency relationship, an incremental frequency stimulation protocol was used: 0.2 ms pulses at 10, 25, 40, 60, 80, 100, 125, and 150 Hz with 1 min in between stimulations.

### Echocardiography

To determine the effect of GnP on cardiac mass and function, we performed echocardiography using the Vevo® 2100 system (Fujifilm VisualSonics Inc., Toronto, Canada). The mice were anesthetized using isoflurane to assess the cardiac function and cardiac muscle mass at the heart rate of 400-500 beats per minute. The measurements of left ventricular mass (LV), ejection fraction (EF), fractional shortening (FS), LV internal diameter (diastole/systole) (LVIDd/s) and posterior wall thickness (diastole/ systole) (PWTd/s) were obtained using the M-mode scanning of the left ventricular chamber.

### Dystrophin staining

The quadriceps muscle was fixed using mounting media and isopentane and 10-µm cryosections sections were used for immunostaining. The sections were washed with PBS and incubated with blocking buffer for 1 hr. at RT. The sections were washed with PBS and incubated with primary antibody (1:100; MANDRA11; DSHB) overnight at 4°C, followed by PBS washes. A goat anti-mouse AlexaFluor 594 florescent secondary antibody was used at a 1:1000 dilution (A-11032, Thermo Fisher; Waltham, MA, USA) for 1 hr. at RT. After another PBS wash, DAPI was added and incubated for 2-5 minutes followed by a PBS wash and coverslips were placed after adding Prolong Gold Antifade mounting media.

### RNA isolation, library preparation and Sequencing

Total RNA was isolated from skeletal and cardiac muscle using the miRNeasy kit (217004, Qiagen). Quadriceps muscle and the apex of the heart were used for total RNA. Total RNA samples were first evaluated for their quantity and quality using TapeStation (Agilent). All the samples were good quality with RIN (RNA Integrity Number) greater than 7. One hundred nanograms of total RNA was used for library preparation with the KAPA mRNA Hyperprep Kit (KK8581) (Roche). Each resulting in a uniquely dual-indexed library was quantified and quality accessed by Qubit and TapeStation. The libraries pooled in equal molarity were sequenced with 2×100bp paired-end configuration on an Illumina NovaSeq 6000 sequencer using the v1.0 reagent kit.

The sequencing reads generated were first quality checked using FastQC (v.0.11.5, Babraham Bioinformatics, Cambridge, UK). The sequence data were next mapped to the mouse reference genome mm10 using the RNA-seq aligner STAR (v.2.5) with the following parameter: “--outSAMmapqUnique 60”. To evaluate quality of the RNA-seq data, the number of reads that fell into different annotated regions (exonic, intronic, splicing junction, intergenic, promoter, UTR, etc.) of the reference genome was assessed using bamutils (from ngsutils v.0.5.9). Uniquely mapped reads were used to quantify the gene level expression employing featureCounts (subread v.1.5.1) with the following parameters: “-s 2-Q 10”. The data was normalized using TMM (trimmed mean of M values) method. Differential expression analysis was performed using edgeR (v.3.12.1). Differential expressed genes were identified at 1.5-fold change and FDR of 0.05 using edgeR^16^. Pathway analysis was performed using Ingenuity Pathway analysis and Metascape^19^.

### Protein Preparation from Skeletal Muscle Tissue

Approximately 30-50 mg of flash frozen quadriceps skeletal muscle tissue was homogenized in 8 M urea, 100mM Tris hydrochloride, pH 8.0 (CHEBI: 16199; CHEBI: 9754) lysis buffer using a BeadBugTM 6 bead mill homogenizer (Benchmark scientific Cat No: D1036), with Precellys 2.8mm stainless steel metal beads (Bertin Corp Cat No: P000925-LYSKO-A) for 10 cycles of 30 second pulses set at 4350 revolutions per minute (RPM) at 4°C. The homogenates were centrifuged (14,000 g x 20 min) to pellet cellular debris and the collected supernatant was sonicated on high power for 20 cycles each of 30 second on/off pulses using a Bioruptor® sonication system (Diagenode Inc. USA, North America cat number B01020001) at 4°C. After sonication, the lysates were centrifuged (14,000 g x 20 min), and the clarified supernatants were quantified using a Bradford protein assay (BioRad Cat No: 5000006).

### Protein Preparation from Cardiac Muscle Tissue

Myocardium proteins were extracted from cryo-fractured tissue using an automated tissue pulverizer (cyroPREP Tissue Disruption System, Covaris model CP02) prior to the addition of urea lysis buffer and sonication.

### Protein Reduction, Alkylation, and Digestion

Following either extraction method, 80 µg of equivalent proteins from sample were treated with 5mM tris (2-carboxyethyl) phosphine hydrochloride (Sigma-Aldrich Cat No: C4706) for 15 minutes at 37°C to reduce the disulfide bonds. The resulting free cysteine thiols were alkylated using 10 mM choloracetamide (Sigma Aldrich Cat No: C0267) for 15 minutes at RT, protected from light. Proteolytic digestion was carried out with Trypsin/LysC Gold (Mass Spectrometry grade, Promega Corporation Cat No: V5072) at an enzyme to protein ratio of 1:40. The two-step enzymatic proteolysis first incubated all study samples for 4 hours at 37°C before diluting the urea concentration to a final concentration of 2 M with 100 mM Tris.HCl (Sigma-Aldrich Cat No: 10812846001) and continued digestion overnight at room temperature.

### Tandem Mass Tag Isobaric Labeling

Digestions were quenched 0.4% trifluoroacetic acid (v/v, Fluka Cat No: 91699) and the resultant peptides were desalted by solid phase extraction using Sep-Pak® Vac cartridges C18 cartridges (WatersTM Cat No: WAT054955), lyophilized O/N, and resuspended in 55 µL of 50 mM triethylammonium bicarbonate (TEAB, Sigma-Aldrich Cat No: T7408), pH 8.5. Peptides were quantified using Quantitative Colorimetric Peptide Assay (Pierce Cat No: 23275) to ensure equivalent concentrations across each set of samples before being covalently labeled with TMTpro™ Isobaric Label Reagent 16-plex (Thermo Fisher Scientific Cat No: 44520, Lot VE299609 or VJ316536 Table #) at a 1:7 peptide to TMTpro ratio. After a 1 hr. incubation, the labeling reaction was quenched with 0.3% hydroxylamine (v/v) for 15 minutes before combining the samples. The multiplexed sample was concentrated to dryness in vacuum centrifuge, reconstituted with 0.1% TFA aq. (v/v), desalted via Waters Sep-Pak® Vac cartridges as before, and lyophilized.

### Identification of differentially expressed proteins

Spectral data for each peptide were analyzed by Proteome DiscovererTM 2.4 service pack. A database of Mus musculus proteins were downloaded from the Universal Protein Knowledgebase/Translated European Molecular Biology Laboratory (UniProtKB/TrEMBL; UP000000589, 55,366 proteins, last modified March 07, 2021) and supplemented with frequently observed contaminants. For the SEQUEST HT search, precursor mass tolerance was set to 10 ppm and fragment mass tolerance set at 0.02 Da. Dynamic modifications include methionine oxidation; phosphorylation on the serine, threonine, and tyrosine residues; as well as lysine modification of acetylation and TMTpro labeling. Static modifications were listed to identify carbamidomethylation on cysteines along with amino terminal TMTpro labeling. For the Proteome Discoverer consensus workflow, isobaric impurities were accounted for, the reporter ion co-isolation threshold was set to 50%, and the average signal to noise threshold was 7. Percolator false discovery rate (FDR) filtration of 1% was applied to both the peptide-spectrum match and protein levels. Differential expression of the protein groups was assessed by ANOVA calculated p-values. Differentially expressed proteins were subjected to pathway analysis using Metascape tool^19^.

### Bone morphometry analysis

Micro-computed tomography analysis was performed using the femur bones on all three groups using a high-resolution desktop _μ_CT (_μ_CT-35; Scano Medical AG, Basel, Switzerland). The bones were scanned at 10 _μ_m resolution, 55kVp intensity with 0.5 mm aluminum filter, and 400 ms integration time. To identify a consistent reconstruction region, femur lengths were measured prior to scanning. The medullary total volume (TV), trabecular bone volume (BV), trabecular thickness (Tb. Th), trabecular number (Tb. N), trabecular separation (Tb. Sp), and structural model index (SMI) cortical bone volume (Ct.BV) and cortical thickness (Ct. Th) at the midshaft region were measured using standard procedures^20^.

## Results

### GnP prevented muscle wasting, muscle function and reduced tumor burden

The overall study design is presented in Figure 1A. Briefly, we had three experimental groups: sham surgery receiving saline injections (control), KPC receiving saline injections (KPC), KPC receiving gemcitabine and nab-paclitaxel (KPC+GnP). Injections occurred on days 4 and 10. Body weights were recorded on alternate days; muscle function test was performed the day before euthanasia (day 13) and echocardiography for cardiac function tests were performed on the day of euthanasia (day 14).

**Figure 1:**
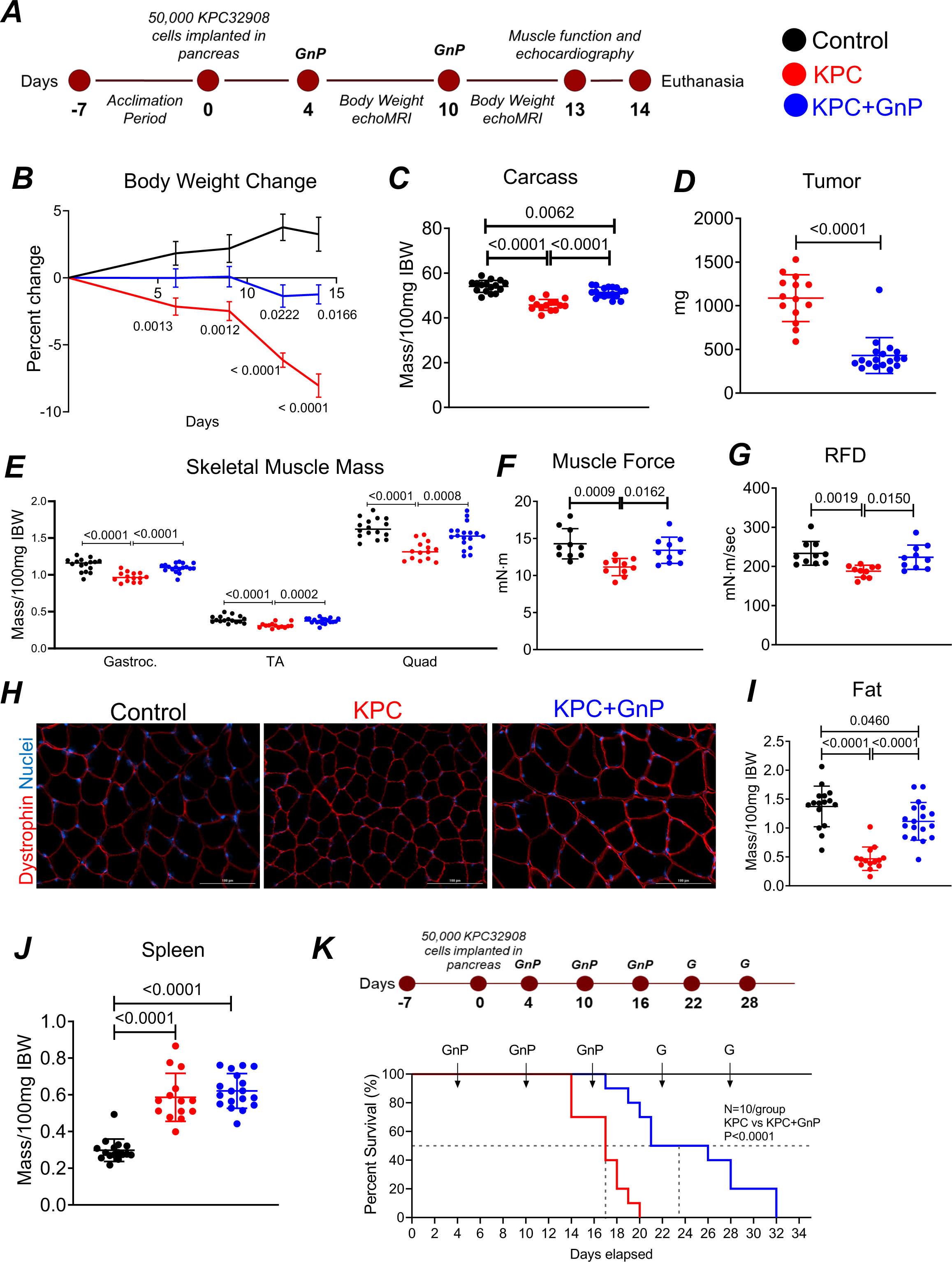
GnP preserved muscle mass and function and reduced tumor burden. (A) Overall experimental plan, 12-week-old male C57BL/6J mice was used in these studies. (B) Percent change in body weight over time until the experimental end point. Statistical values compare timepoint within group to pre. One-way repeated measures ANOVA. (C) Carcass weights. (D) Tumor weights. (E) Muscle weights normalized to 100mg initial body weight. (F,G) in vivo muscle contractility test. RFD: Rate of force development. (H) Representative images of gastrocnemius muscle stained for dystrophin for cross sectional area assessment. (I, J) Epididymal fat and spleen weights normalized to 100mg initial body weights. Data are expressed as mean ± standard deviation. (B-J: n=12-15 per group). (C-G, I, J: One-way ANOVA statistical test were used). (K) Kaplan-Meier survival estimate. The dashed line represents median survival, n =10 per group.

Body weight at the end of the study decreased 8% in KPC compared to control, and KPC+GnP had less than 3% weight loss which was also reflected in the carcass (Figure 1B). Carcass weight was decreased in KPC compared to control and KPC+GnP (Figure 1C). At the end of the study, tissues were collected and weighed. Additionally, GnP administration halved tumor burden when compared to KPC (Figure 1D, p<0.0001). There was a 14%, 18% and 20% loss of gastrocnemius (p<0.0001), tibialis anterior (p<0.0001) and quadriceps (p<0.0001) muscles, respectively in KPC group when compared against controls (Figures 1E). KPC+GnP prevented muscle mass loss compared to KPC. KPC+ GnP muscle mass weight was similar to controls. KPC exhibited decreased force (Figure 1F) and rate of force development (Figure 1G) compared to control and KPC+GnP. No significant differences were observed in force and rate of force development of KPC+GnP when compared to controls, indicating muscle dysfunction and weakness was preserved with GnP. By dystrophin staining, we observe that KPC+GnP preserved muscle size compared to KPC (Figure 1H). KPC had a 70% reduction in fat mass, which was partially attenuated in KPC+GnP, when compared against controls (Figure 1I). Splenomegaly was observed in both KPC and KPC+GnP (Figure 1J). These results indicate that GnP treatment can preserve both muscle mass and function in an experimental model of PDAC cachexia.

Given these pronounced effects of GnP on tumor burden and indices of cachexia, we determined if GnP extended survival in our experimental model of PDAC cachexia. GnP increased survival, with a median survival of 23.5 d in KPC+GnP compared with 17 d in KPC (Figure 1K, p=0.0002).

### Skeletal muscle RNA sequencing analysis shows that GnP treatment preserved several pathways involved in muscle wasting

Principal component analysis (PCA) clustered control and KPC group distinctly but showed no separation between control and KPC+GnP (Figure 2A). In skeletal muscle, we found 4944 differentially expressed genes between KPC and controls at 1.5-fold change and at false discovery rate of (FDR) of 0.05 (Supplementary Table 1). There were no differentially expressed genes identified between KPC+GnP and control, reflecting the clustering pattern seen in the PCA. Heatmap showed differentially expressed genes had a distinct direction of effect between controls and KPC and such effect was reduced or reversed in KPC+GnP (Figure 2B). Several well-known muscle wasting pathways were identified including oxidative phosphorylation, phospholipase C signaling, hedgehog signaling, calcium signaling, sirtuin signaling (Figure 2C). Genes identified in these pathways were dysregulated in KPC but were either reversed or reduced in KPC+GnP (Figure 2D). The complete list of pathways between KPC and control is presented in Supplementary Table 2.

**Figure 2:**
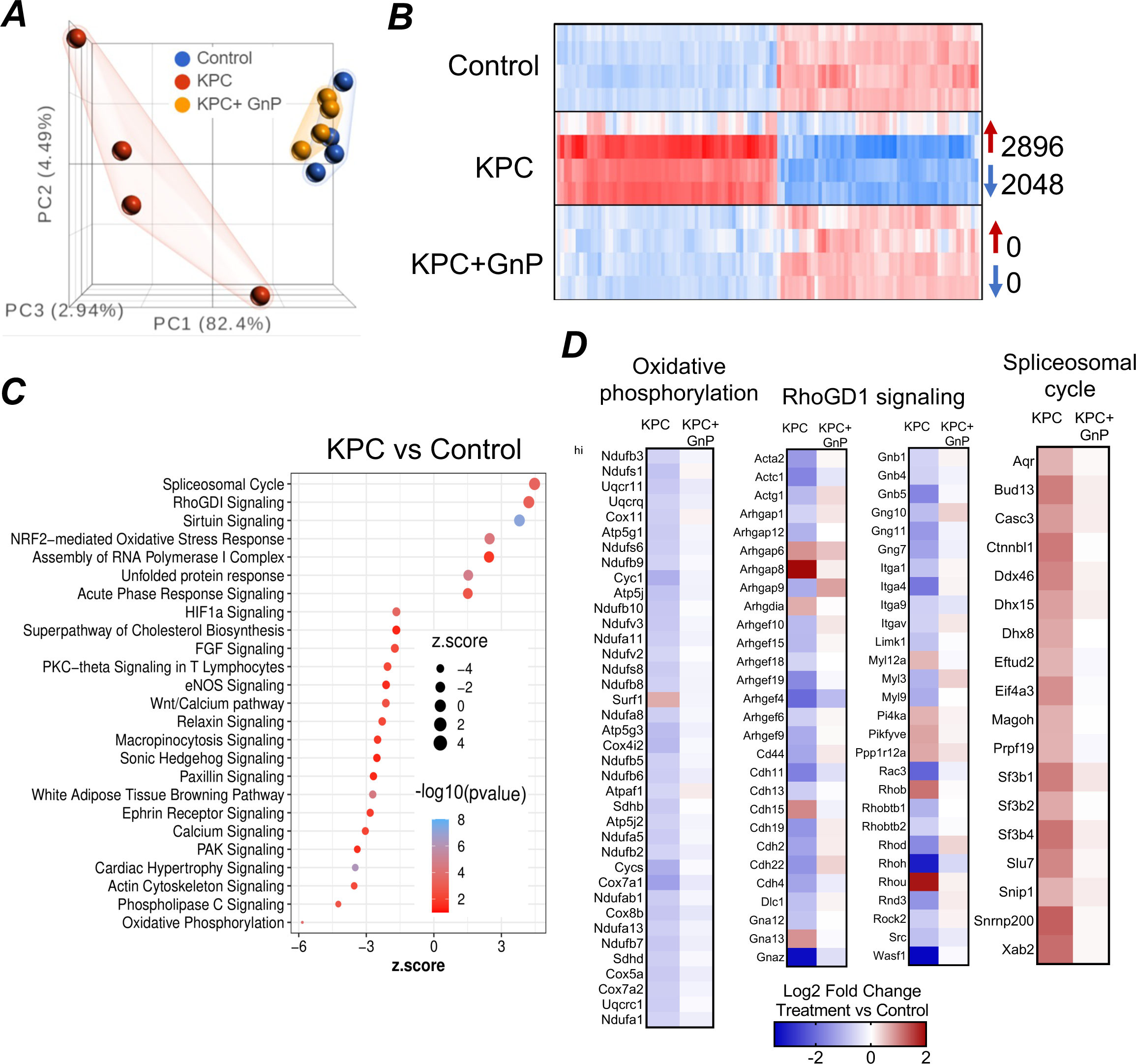
GnP treatment reduced the expression of genes associated with muscle wasting and inflammatory pathways in skeletal muscle. (A) Principal component analysis shows separation between control and KPC group while clustering with KPC+GnP (n=4 per group). (B) Heatmap of differentially expressed genes (1.5-fold change, p< 0.05) with red and blue color indicating up and downregulation of genes, respectively. (C) Representative pathways using differentially expressed genes (4944 genes) from KPC vs control comparison. (D) Representative pathways showing that GnP decreased or reversed the expression of genes involved in muscle wasting and weakness.

To recapitulate some of the previous findings, we checked for the expression of several well studied cachexia genes such as Trim63, Fbxo32, Lc3b, Atf4, Foxo1, Ucp2, Ucp3 and Pdk4. All of these genes are upregulated in an experimental model of PDAC cachexia^16^. Similar results were observed in the current study. The expression of these genes was reversed in KPC+GnP. These results suggest that GnP treatment may effectively prevent perturbations to muscle wasting pathways previously reported in experimental models of PDAC cachexia.

### Skeletal muscle proteomics analysis revealed that GnP treatment reduced protein expression associated with muscle metabolism and inflammation

Gastrocnemius muscle was used for proteome profiling. There were 228 and 66 differentially expressed proteins identified for KPC vs control and KPC+GnP vs control, respectively (Figure 3A). Several pathways associated with muscle metabolism such as citric acid cycle, pyruvate metabolism and mitochondrial translation were identified (Figure 3B). The TCA cycle and electron transportation chain was distinctly associated to KPC+GnP vs control. The expression of proteins associated with these pathways were either reduced or reversed in KPC+GnP (Figure 3C). As well, proteins in several inflammatory pathways, including TNF_α_/NF-_κ_B and platelet signaling (Figure 3C) were reduced in KPC+GnP, indicating skeletal muscle has reduced protein inflammation. When we overlapped RNA Seq and proteomics data, 80 molecules were altered in the same direction (Figure 3D, Supplemental Figure 1A and Supplemental Table 7). To corroborate with previous findings, we found several cachexia related genes such as Trim63, Ei4ebp1, Pdk4, Fkbp5, Tfrc to have similar direction of effect in RNA seq and proteomics data, which can be considered as a validation at the protein level. In all, these results suggest that GnP treatment preserved disruptions to the muscle metabolism pathway and inflammation in experimental models of PDAC cachexia. The list of differentially expressed proteins is presented in Supplementary Table 3.

**Figure 3:**
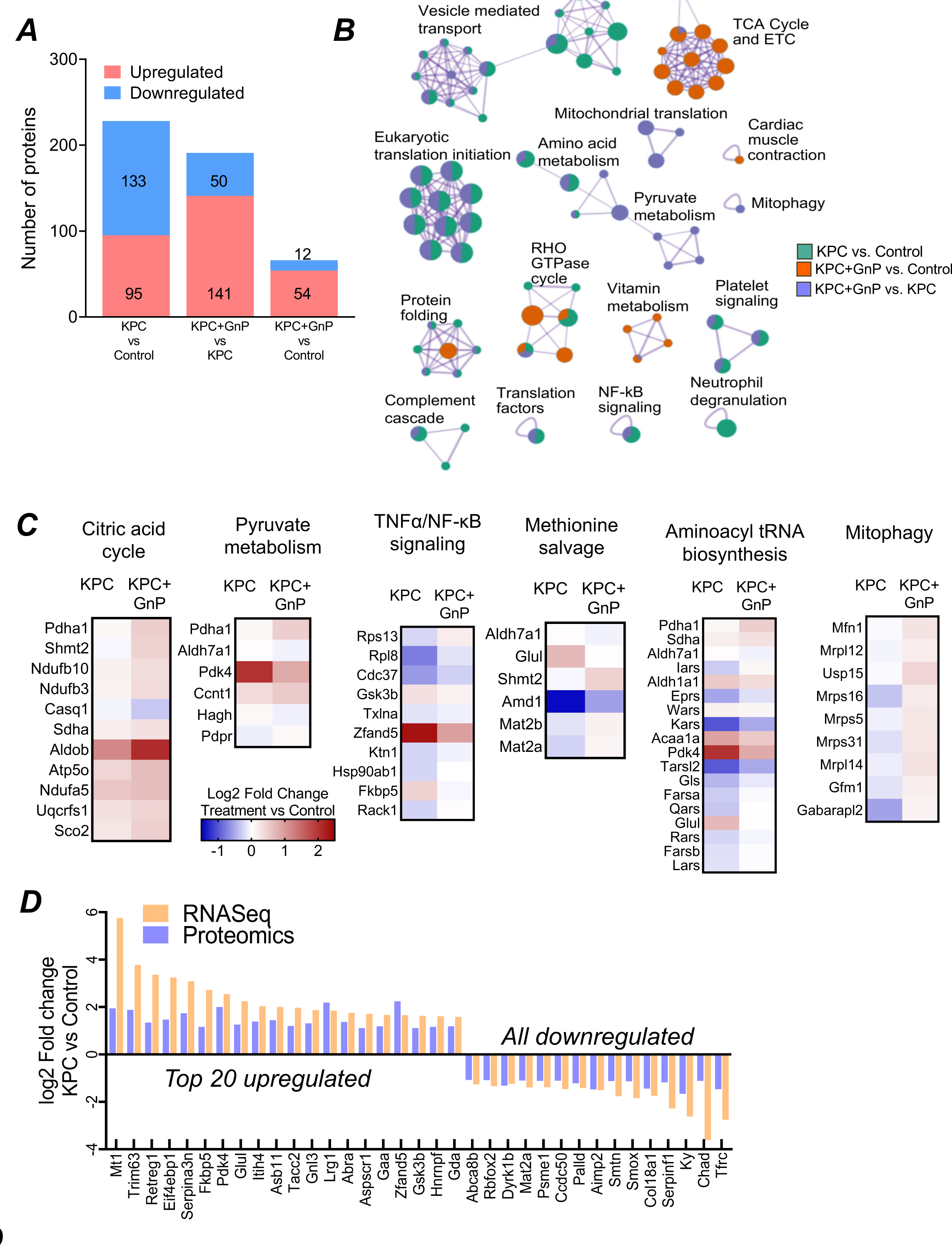
GnP treatment altered protein expression associated with muscle metabolism and inflammation. (A) Differentially expressed proteins in different comparisons with pink and blue indicating up and downregulated proteins, respectively (n=4 per group). (B) Pathways identified using differentially expressed proteins from (A) using Metascape. Pathways shared between groups are indicated with shared colors. (C) GnP treatment reduced or reversed the expression of proteins involved in inflammation and muscle metabolism. (D) Differentially expressed genes and proteins were overlapped to identify common features. The top 20 upregulated molecules and all of the downregulated molecules are presented.

### GnP treatment preserved cardiac mass and function

While the effect of chemotherapy on muscle wasting has been well studied, its effect on cardiac function remains less explored in the context of pancreatic cancer cachexia. Therefore, we defined the effect of KPC and GnP treatment on cardiac muscle mass and function. Cardiac mass, measured by wet tissue weight, was reduced in KPC compared to control (p<0.0001) and KPC+GnP (p=0.0012). There were no differences between KPC+GnP and control (Figure 4A). In addition, there was a decrease in left ventricular (LV) mass, measured by echocardiography, in KPC compared to KPC+GnP (p=0.0191) and control (p=0.0151, Figure 4B). No difference was observed in LV mass between KPC+GnP and control. Despite a decrease in LV and whole cardiac mass in KPC, the ejection fraction (p<0.001, Figure 4C) and fractional shortening (p=0.0001, Figure 4D) was increased in KPC relative to the control.

**Figure 4:**
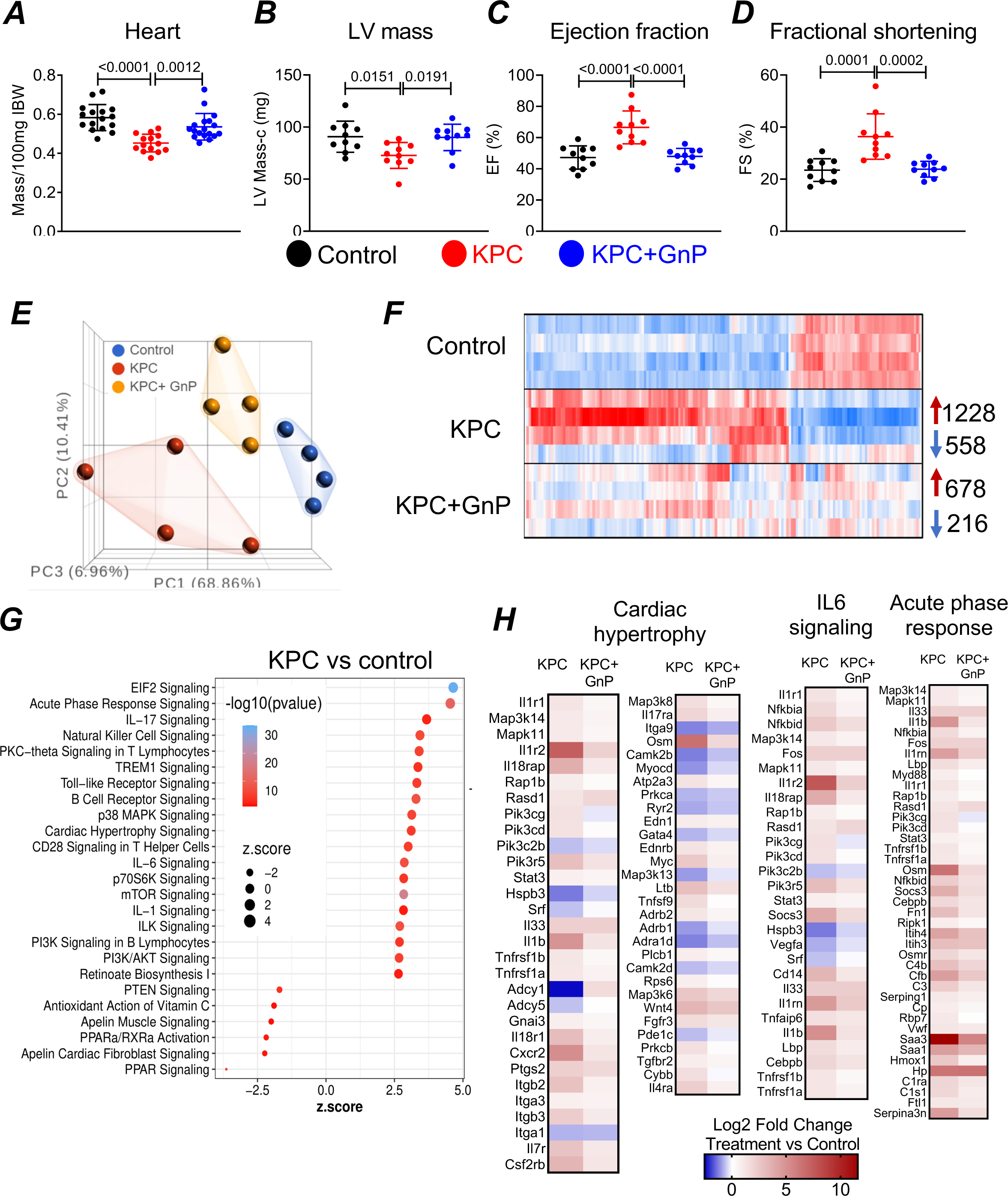
GnP treatment preserved cardiac mass and function. (A) Cardiac mass normalized to 100mg initial body weight. (B-D) Left ventricular, ejection fraction and fractional shortening were measured using echocardiography (n=8-20 per group). (E) Principal component analysis shows separation between all three groups (n=4 per group). (F) Heatmap of differentially expressed genes (1.5-fold change, p< 0.05) with red and blue color indicating up and downregulation of genes. (G) Representative pathways using differentially expressed genes (1786 genes) from KPC vs control comparison. (H) Representative pathways showing that GnP decreased or reversed the expression of genes involved in cardiac dysfunction.

RNA Seq on flash frozen heart tissue was performed. PCA identified three clusters representing the three groups: control, KPC and KPC+GnP (Figure 4E). While the heatmap shows a distinct expression pattern between KPC and control, roughly half of these genes were reduced or rescued in KPC+GnP (Figure 4F). KPC vs control had 1786 and KPC+GnP vs control had 894 differentially expressed genes identified (Supplementary Table 4). Several inflammatory signaling, acute phase response, PI3K signaling and cardiac hypertrophy signaling pathways were identified (Figure 4G). KPC+GnP reduced the expression of the genes involved in these pathways, indicating that GnP treatment may reduce inflammation and inhibit signaling pathways involved in cardiac wasting in mice (Figure 4H). The complete list of pathways between KPC and control in heart is presented in Supplementary Table 5.

### Proteomic analysis of cardiac muscle showed that GnP treatment reduced inflammation and modulates lipid metabolism

In cardiac muscle, protein analysis identified 638 and 66 differentially expressed proteins between KPC vs control and KPC+GnP vs control, respectively (Figure 5A). Pathway analysis identified several pathways associated with lipid metabolism including cholesterol metabolism, chylomicron assembly and fatty acid metabolism (Figure 5B) which have been linked to several heart diseases. The majority of proteins expressed in these pathways are upregulated in the KPC vs control comparison and it is known that increased lipoprotein levels are considered as risk factor for cardiovascular disease. The expression of these proteins was reduced in KPC+GnP vs controls suggesting that GnP may potentially improve cardiac metabolism (Figure 5C). As well, GnP treatment reduced protein levels of inflammatory pathways such as neutrophil degradation (Figure 5C). When we overlapped RNA Seq and proteomics data, 99 molecules were differentially regulated in the same direction (Figure 5D and Supplemental Figure 1). In all, the data suggests that GnP treatment can potentially reduce inflammation and modulates lipid metabolism in the heart. The list of differentially expressed proteins is presented in Supplementary Table 6.

**Figure 5:**
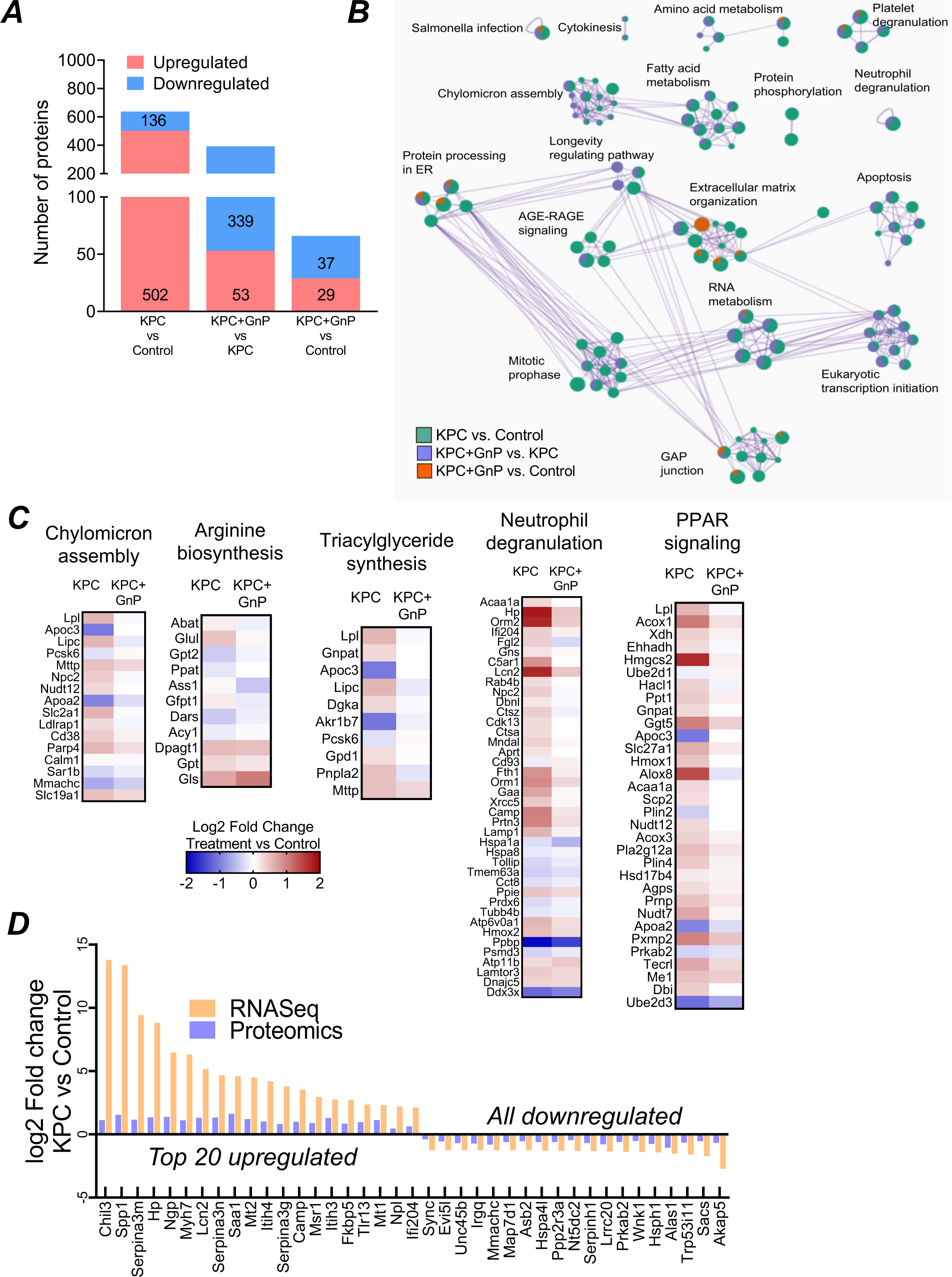
Proteomic analysis of cardiac muscle showed that GnP treatment reduced inflammation and modulates lipid metabolism. (A) Differentially expressed proteins in different comparisons with pink and blue indicating up and downregulated proteins, respectively (n=4 per group). (B) Pathways identified using differentially expressed proteins from (A) using Metascape. Pathways that are shared between groups are indicated with shared colors. (C) GnP treatment reduced or reversed the expression of proteins involved in inflammation and lipid metabolism. (D) Differentially expressed genes and proteins were overlapped to identify common features. The top 20 upregulated molecules and all of the downregulated molecules are presented.

### GnP treatment restored the complete blood counts indicating reduced inflammation

Complete blood count was measured using Heska Element HT5 (Loveland, CO USA). KPC had a significant increase in neutrophils and monocytes compared to control and were reduced in KPC+GnP (Supplemental Figure 2B, D). We calculated the neutrophil/lymphocyte ratio, a higher score is generally associated with poor outcomes. The ratio was increased in KPC compared to control and KPC+GnP, while there was no difference between KPC+GnP versus control (Supplemental Figure 2E). Similar patterns in KPC compared to controls were observed in basophils, eosinophils, and platelets (Supplemental Figure 2F-H). The mean corpuscular volume and mean corpuscular hemoglobin was slightly reduced in KPC+GnP, which may potentially indicate iron deficiency anemia (Supplemental Figure 2I, J).

### Bone alterations were not observed in our experimental model of PDAC Cachexia

There were no observed changes in BV/TV, Tb.Th, Tb.Sp and Tb.N between any groups (Supplemental Figure 2K-N). However, the bone connectivity density was reduced in KPC+ GnP group but not in the KPC group.

## Discussion

It is well documented that GnP alone or in combination with other therapeutic targets improved survival in patients and mice with PDAC^21–23^. However, as most of the patients with pancreatic cancer experience cachexia, it is imperative to have an understanding of systemic GnP effects. Herein, we evaluated the effect of GnP in an experimental model of PDAC cachexia. In line with previous studies, the tumor burden was reduced in GnP treated KPC mice^22^. To the best of our knowledge, this is the first study to show that GnP significantly prevented muscle and cardiac wasting and dysfunction. These findings demonstrate that GnP preserves muscle and heart in mice with aggressive pancreatic cancer cachexia.

The mice in the KPC group experienced pronounced loss of body weight accompanied by skeletal muscle atrophy and weakness and fat loss. RNA sequencing identified atrophy genes such as Trim63/Murf1 and Fbxo32/Atrogin-1^16, 24, 25^. Other genes which are often reported to be upregulated in experimental models of cachexia such as Pdk4 and Ucp2 were also identified in our study^26, 27^. Several well-known cachexia pathways such as oxidative phosphorylation, acute phase response, and sonic hedgehog signaling were dysregulated in KPC group indicating muscle wasting^24, 28^. However, when compared against the controls, several of these pathways were prevented in KPC+GnP group where the muscle wasting was not observed, however, there was a slight decrease in overall body weight towards the end of the study. As expected, skeletal muscle function was reduced in KPC, there was no significant differences between control and GnP, indicating that GnP protects muscle function. In line with the functional data, the expression of genes dysregulated in various pathways associated with muscle wasting and function were either reversed or greatly reduced in the GnP group. Several metabolic changes including alterations in mitochondrial function and TCA cycle were identified and reported in other experimental models^27, 29^. Deep proteome analysis identified several dysregulated proteins including Pdk4, Ndufb10, Ndufb3 and Ndufa5 that play a role in the citric acid cycle and pyruvate metabolism in the KPC group compared to controls. Similar to the transcriptomics data, the expression of these dysregulated proteins was also reduced or reversed in KPC+GnP group at the protein level. As well, some of the well-established cachexia genes including Trim63, Pdk4, Tfrc, Glul, Fkbp5, Mt1 had similar direction of expression in both RNA Seq and proteomics data, providing an additional layer of validation. These findings suggest that GnP treatment preserves skeletal muscle, preventing wasting, weakness and maintaining homeostatic muscle signaling in an experimental model of PDAC cachexia.

Over the course of disease progression, patients with cancer induced cachexia experience cardiac atrophy. A retrospective study in 177 cancer patients, including 60 with pancreatic cancer, demonstrated that patients with cachexia as demonstrated by body weight loss was also associated with decreased heart weight^30^. In a similar study, patients with non-small cell lung cancer were found to experience left ventricular mass atrophy^31^. A significant impairment of cardiac function has been observed in a metastatic colorectal cancer model of cachexia^18^. Similar to these observations, our experimental model of PDAC exhibited decreased left ventricular mass and impaired function in the KPC group. In our study, RNA Seq analysis from the heart identified pathways associated with inflammation, including IL-6 signaling, acute phase response and cardiac hypertrophy. Several signaling pathways, including mTOR and PI3K/Akt signaling pathways were also identified. While their role in skeletal wasting is well established, a previous report showed decreased cardiac protein synthesis accompanied by decreased mTORC1 activity in colorectal cancer model^32^. In our study, proteomics analysis of cardiac muscle identified several pathways associated with lipid metabolism including chylomicron assembly, triglyceride synthesis and fatty acid metabolism. Increased triglyceride levels are a known risk factor of cardiovascular disease. Several proteins including Lpl, Lipc were upregulated in cardiac muscle from KPC mice. Studies have reported that increased cardiac Lpl may lead to abnormal cardiac lipid accumulation eventually, leading to cardiac dysfunction^33^. The expression of proteins involved in cardiac dysfunction were reduced or reversed in KPC+GnP group indicating that GnP may have a protective effect on cardiac function, coinciding with no significant difference in cardiac mass or function in KPC+GnP group. These results indicate that along with preventing the skeletal muscle mass and function, GnP treatment also protects cardiac function in an experimental model of PDAC cachexia.

As inflammation is commonly observed in cancer and cachexia, several inflammatory markers have been measured to understand the disease progression. In particular, an increase in the neutrophil to lymphocyte ratio has been vastly associated with poor prognosis in different cancer types including PDAC^15, 34, 35^ and has been shown to be an independent prognostic factor of progression free overall survival. In line with these observations in several human studies, there was an increase in neutrophil to lymphocyte ratio in KPC when compared to controls. As well, we observed an elevated monocytes, basophils, eosinophils and platelets in KPC compared to control and KPC+GnP. There were no differences between KPC+GnP and controls indicating that GnP treatment may reduce inflammation leading to restored blood counts. No significant differences were observed in several bone parameters in KPC nor KPC+GnP. While osteopenia has been observed in patients with PDAC and associated with reduced survival, we did not observe this in our experimental model of cachexia. A plausible explanation is that bone turnover may require a longer time to exhibit an altered phenotype, as our study lasted only 14 days. This can be studied in future by using a less aggressive pancreatic cancer cell line to understand if there are any bone alterations in PDAC cachexia.

It is well documented that chemotherapy related toxicities lead to substantial weight and muscle loss in patients with cancer and eventually leading cachexia^36, 37^. Similar observations haven been made in experimental models where chemotherapy has been shown to induce side effects resulting in cachexia. For example, administration of Folfiri in experimental models of colon adenocarcinoma induced severe muscle and adipose wasting along with other metabolic derangements^8^. Carboplatin treatment in breast cancer bone metastasis model led to muscle weakness and wasting^38^. To put the current study in this context, GnP prevented skeletal and cardiac muscle wasting and weakness, partially prevented fat wasting and improved survival suggesting that GnP treatment may have a beneficial impact in PDAC cachexia. More clinical studies are needed in the future to understand the impact of GnP on cachexia in patients with PDAC. While different toxicities have been reported with GnP administration including pulmonary toxicity, appropriate interventions led to complete recovery from these toxicities^39^. Herein we show that GnP increased survival in a murine model of PDAC cachexia which is in agreement with previous studies showing improved overall survival in patients with pancreatic adenocarcinoma^6^, a comprehensive investigation is needed to understand if the improved survival is because of reduced cachexia.

In all, our study shows for the first time that gemcitabine and nab-paclitaxel may have protective effects on cachexia. Future clinical trials in pancreatic cancer might consider the addition of cachexia-related endpoints to better understand the translatable nature of these findings.

---

**Supplemental Figure 1:**
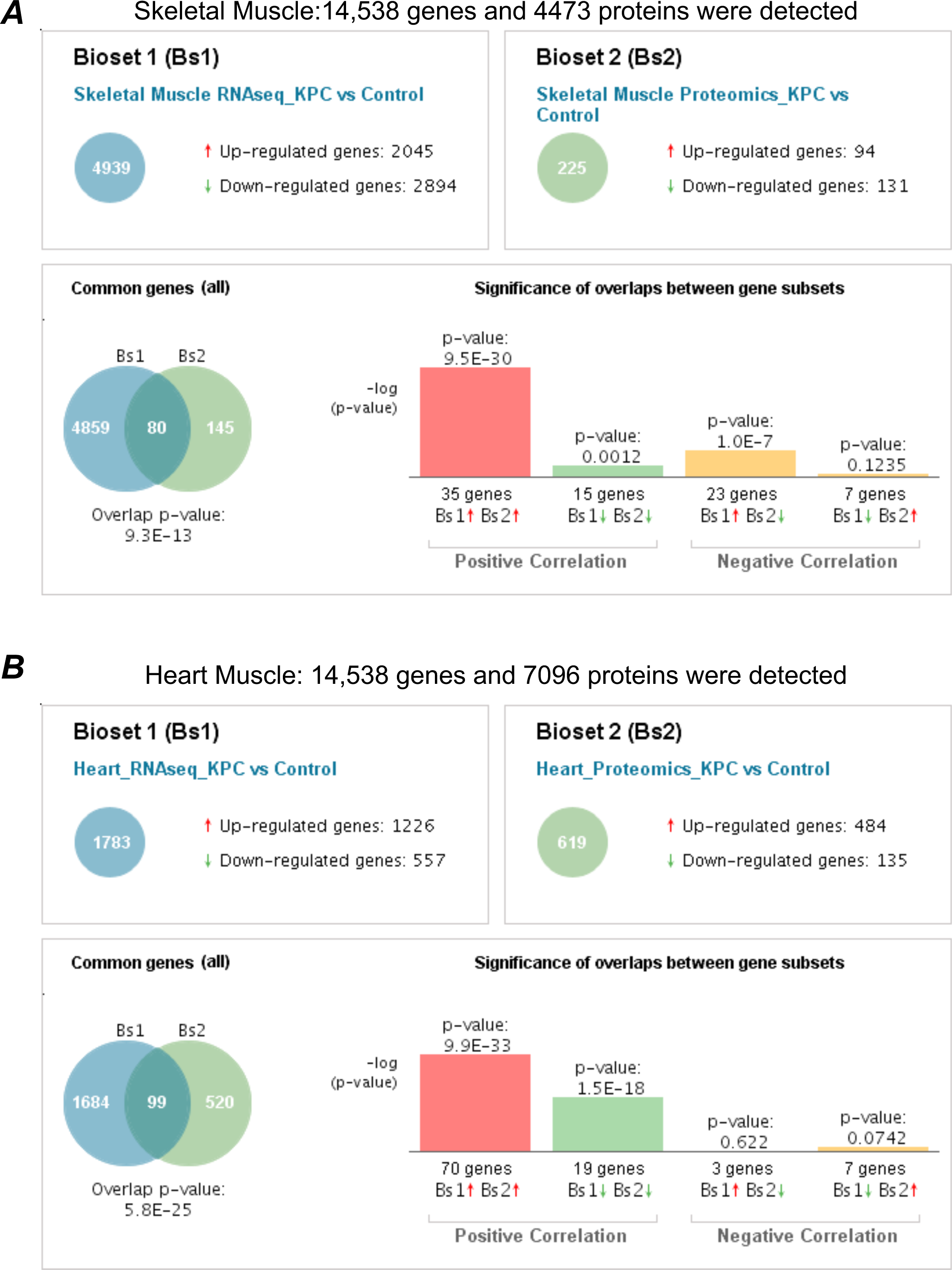
Differentially expressed genes and proteins. (A) Differentially expressed genes (blue) and proteins (green) correlated in skeletal muscle. RNA seq identified 14, 538 genes and 4,473 proteins were identified in the deep proteome profiling. (B) Differentially expressed genes (blue) and proteins (green) correlated in cardiac muscle. RNA seq identified 14, 538 genes and 7,096 proteins were identified in the deep proteome profiling.

**Supplementary Figure 2:**
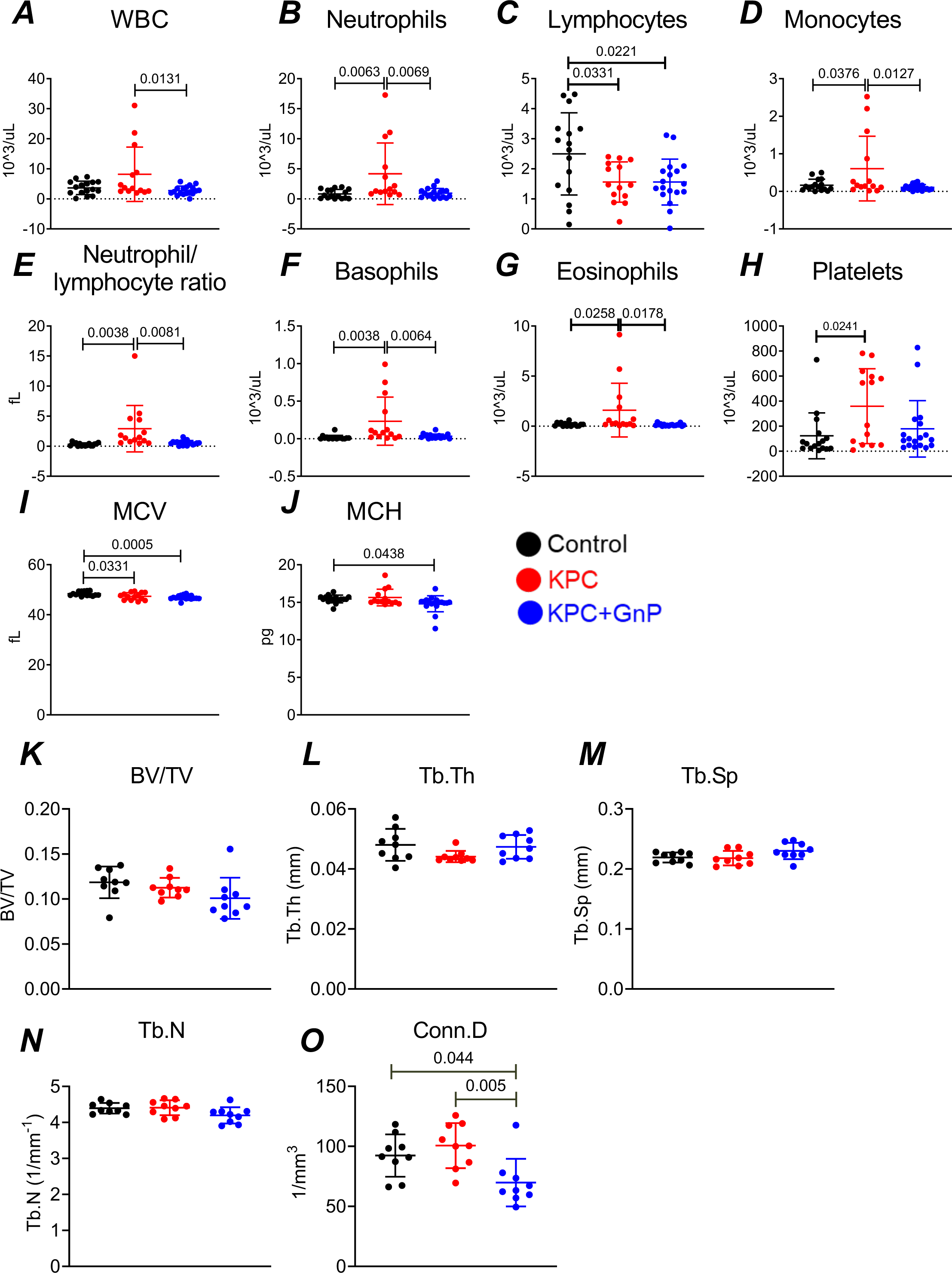
GnP treatment restored the blood counts and no significant bone alterations. (A-J) Complete blood count parameters shown along with significant difference (n=12-15 per group). (K-L) Bone morphometry results showed no significant difference except for trabecular connectivity density. Abbreviations: (I’m not familiar with these abbreviations for bone markers).

**Supplementary Table S1:** List of differentially expressed genes identified from skeletal muscle using RNA Seq (related to Figure 2).

**Supplementary Table S2:** List of significant pathways generated using differentially expressed genes from skeletal muscle from the KPC versus sham comparison (related to figure 2).

**Supplementary Table S3:** Differentially expressed proteins identified in skeletal muscle (related to Figure 3).

**Supplementary Table S4:** List of differentially expressed genes identified from cardiac muscle using RNA Seq (related to Figure 4).

**Supplementary Table S5:** List of significant pathways generated using differentially expressed genes from cardiac muscle identified from KPC versus sham comparison (related to Figure 4).

**Supplementary Table S6:** Differentially expressed proteins identified in cardiac muscle (related to Figure 5).

**Supplementary Table S7:** List of differentially expressed genes and proteins in skeletal and cardiac muscle (related to Figures 3,5 and Supplemental Figure 1).

## References

1. Hendifar AE, Chang JI, Huang BZ, et al. Cachexia, and not obesity, prior to pancreatic cancer diagnosis worsens survival and is negated by chemotherapy. J Gastrointest Oncol 2018;9:17–23.

2. Nemer L, Krishna SG, Shah ZK, et al. Predictors of Pancreatic Cancer-Associated Weight Loss and Nutritional Interventions. Pancreas 2017;46:1152–1157.

3. Naumann P, Eberlein J, Farnia B, et al. Continued Weight Loss and Sarcopenia Predict Poor Outcomes in Locally Advanced Pancreatic Cancer Treated with Chemoradiation. Cancers 2019;11:709.

4. Damrauer JS, Stadler ME, Acharyya S, et al. Chemotherapy-induced muscle wasting: association with NF-_κ_B and cancer cachexia. Eur J Transl Myol 2018;28:7590.

5. Conroy T, Desseigne F, Ychou M, et al. FOLFIRINOX versus gemcitabine for metastatic pancreatic cancer. N Engl J Med 2011;364:1817–1825.

6. Von Hoff DD, Ervin T, Arena FP, et al. Increased survival in pancreatic cancer with nab-paclitaxel plus gemcitabine. N Engl J Med 2013;369:1691–1703.

7. Sorensen JC, Cheregi BD, Timpani CA, et al. Mitochondria: Inadvertent targets in chemotherapy-induced skeletal muscle toxicity and wasting? Cancer Chemother Pharmacol 2016;78:673–683.

8. Pin F, Barreto R, Couch ME, et al. Cachexia induced by cancer and chemotherapy yield distinct perturbations to energy metabolism. J Cachexia Sarcopenia Muscle 2019;10:140– 154.

9. Barreto R, Waning DL, Gao H, et al. Chemotherapy-related cachexia is associated with mitochondrial depletion and the activation of ERK1/2 and p38 MAPKs. Oncotarget 2016;7:43442–43460.

10. Petrillo A, Pappalardo A, Pompella L, et al. Nab-paclitaxel plus gemcitabine as first line therapy in metastatic pancreatic cancer patients relapsed after gemcitabine adjuvant treatment. Med Oncol Northwood Lond Engl 2019;36:83.

11. Riedl JM, Posch F, Horvath L, et al. Gemcitabine/nab-Paclitaxel versus FOLFIRINOX for palliative first-line treatment of advanced pancreatic cancer: A propensity score analysis. Eur J Cancer Oxf Engl 1990 2021;151:3–13.

12. Burris HA, Moore MJ, Andersen J, et al. Improvements in survival and clinical benefit with gemcitabine as first-line therapy for patients with advanced pancreas cancer: a randomized trial. J Clin Oncol Off J Am Soc Clin Oncol 1997;15:2403–2413.

13. Pusceddu S, Ghidini M, Torchio M, et al. Comparative Effectiveness of Gemcitabine plus Nab-Paclitaxel and FOLFIRINOX in the First-Line Setting of Metastatic Pancreatic Cancer: A Systematic Review and Meta-Analysis. Cancers 2019;11:484.

14. Klein-Brill A, Amar-Farkash S, Lawrence G, et al. Comparison of FOLFIRINOX vs Gemcitabine Plus Nab-Paclitaxel as First-Line Chemotherapy for Metastatic Pancreatic Ductal Adenocarcinoma. JAMA Netw Open 2022;5:e2216199.

15. Suzuki R, Takagi T, Hikichi T, et al. Derived neutrophil/lymphocyte ratio predicts gemcitabine therapy outcome in unresectable pancreatic cancer. Oncol Lett 2016;11:3441–3445.

16. Rupert JE, Narasimhan A, Jengelley DHA, et al. Tumor-derived IL-6 and trans-signaling among tumor, fat, and muscle mediate pancreatic cancer cachexia. J Exp Med 2021;218:e20190450.

17. Huot JR, Pin F, Bonetto A. Muscle weakness caused by cancer and chemotherapy is associated with loss of motor unit connectivity. Am J Cancer Res 2021;11:2990–3001.

18. Huot JR, Pin F, Narasimhan A, et al. ACVR2B antagonism as a countermeasure to multi-organ perturbations in metastatic colorectal cancer cachexia. J Cachexia Sarcopenia Muscle 2020;11:1779–1798.

18. Zhou Y, Zhou B, Pache L, et al. Metascape provides a biologist-oriented resource for the analysis of systems-level datasets. Nat Commun 2019;10:1523.

20. Robling AG, Kang KS, Bullock WA, et al. Sost, independent of the non-coding enhancer ECR5, is required for bone mechanoadaptation. Bone 2016;92:180–188.

21. Kunzmann V, Ramanathan RK, Goldstein D, et al. Tumor Reduction in Primary and Metastatic Pancreatic Cancer Lesions With nab-Paclitaxel and Gemcitabine: An Exploratory Analysis From a Phase 3 Study. Pancreas 2017;46:203–208.

22. McDonald PC, Chafe SC, Brown WS, et al. Regulation of pH by Carbonic Anhydrase 9 Mediates Survival of Pancreatic Cancer Cells With Activated KRAS in Response to Hypoxia. Gastroenterology 2019;157:823–837.

23. Wade SJ, Sahin Z, Piper A-K, et al. Dual Delivery of Gemcitabine and Paclitaxel by Wet-Spun Coaxial Fibers Induces Pancreatic Ductal Adenocarcinoma Cell Death, Reduces Tumor Volume, and Sensitizes Cells to Radiation. Adv Healthc Mater 2020;9:e2001115.

24. Argilés JM, Busquets S, Stemmler B, et al. Cancer cachexia: understanding the molecular basis. Nat Rev Cancer 2014;14:754–762.

25. Wang G, Biswas AK, Ma W, et al. Metastatic cancers promote cachexia through ZIP14 upregulation in skeletal muscle. Nat Med 2018;24:770–781.

26. Busquets S, Almendro V, Barreiro E, et al. Activation of UCPs gene expression in skeletal muscle can be independent on both circulating fatty acids and food intake. Involvement of ROS in a model of mouse cancer cachexia. FEBS Lett 2005;579:717–722.

27. Pin F, Novinger LJ, Huot JR, et al. PDK4 drives metabolic alterations and muscle atrophy in cancer cachexia. FASEB J Off Publ Fed Am Soc Exp Biol 2019;33:7778–7790.

28. Baracos VE, Martin L, Korc M, et al. Cancer-associated cachexia. Nat Rev Dis Primer 2018;4:17105.

29. Brown JL, Rosa-Caldwell ME, Lee DE, et al. Mitochondrial degeneration precedes the development of muscle atrophy in progression of cancer cachexia in tumour-bearing mice. J Cachexia Sarcopenia Muscle 2017;8:926–938.

30. Barkhudaryan A, Scherbakov N, Springer J, et al. Cardiac muscle wasting in individuals with cancer cachexia. ESC Heart Fail 2017;4:458–467.

31. Kazemi-Bajestani SMR, Becher H, Butts C, et al. Rapid atrophy of cardiac left ventricular mass in patients with non-small cell carcinoma of the lung. J Cachexia Sarcopenia Muscle 2019;10:1070–1082.

32. Manne NDPK, Lima M, Enos RT, et al. Altered cardiac muscle mTOR regulation during the progression of cancer cachexia in the ApcMin/+ mouse. Int J Oncol 2013;42:2134–2140.

33. Pulinilkunnil T, Rodrigues B. Cardiac lipoprotein lipase: metabolic basis for diabetic heart disease. Cardiovasc Res 2006;69:329–340.

34. Howard R, Kanetsky PA, Egan KM. Exploring the prognostic value of the neutrophil-to-lymphocyte ratio in cancer. Sci Rep 2019;9:19673.

35. Shibutani M, Maeda K, Nagahara H, et al. A high preoperative neutrophil-to-lymphocyte ratio is associated with poor survival in patients with colorectal cancer. Anticancer Res 2013;33:3291–3294.

36. Silver HJ, Dietrich MS, Murphy BA. Changes in body mass, energy balance, physical function, and inflammatory state in patients with locally advanced head and neck cancer treated with concurrent chemoradiation after low-dose induction chemotherapy. Head Neck 2007;29:893–900.

37. Awad S, Tan BH, Cui H, et al. Marked changes in body composition following neoadjuvant chemotherapy for oesophagogastric cancer. Clin Nutr Edinb Scotl 2012;31:74–77.

38. Hain BA, Xu H, Wilcox JR, et al. Chemotherapy-induced loss of bone and muscle mass in a mouse model of breast cancer bone metastases and cachexia. JCSM Rapid Commun 2019;2:e00075.

39. Baig J, Shokouh-Amiri M, Chan J, et al. The Spectrum of Pulmonary Toxicity in Pancreatic Cancer Patients Receiving Gemcitabine Combination Chemotherapy. Case Rep Oncol 2019;12:506–512.

